# Iodine status of children and Knowledge, Attitude, Practice of Iodised salt use in a remote community in Kerema district, Gulf province, Papua New Guinea

**DOI:** 10.1101/317230

**Authors:** Janny M Goris, Victor J Temple, Nienke Zomerdijk, Karen Codling

**Affiliations:** PNG Foundation; University of Papua New Guinea, School of Medicine and Health Sciences; The University of Queensland, School of Medicine; Regional Coordinator for Southeast Asia and the Pacific, Iodine Global Network

## Abstract

Iodine deficiency is the single most common cause of preventable mental impairment in communities with suboptimal iodine intake. Objective of the present study was to assess in more detail the iodine status and knowledge, attitudes and practice (KAP) relating to use of iodised salt in a remote community in Kotidanga area, Kerema district, Gulf province, Papua New Guinea. This prospective school and community based cross-sectional study was carried out in 2017. Simple random sampling was used to select schools. Multistage sampling was used to randomly select 291 children aged 6 to 12 years. Salt samples were collected for analysis from children’s households as well as a single urine sample of selected children. Salt iodine content and Urinary iodine concentration (UIC) were analysed. A semi-structured FAO questionnaire was used to assess KAP of three different community groups. Only 64% of households had salt on the day of data collection. Mean iodine content in household salt samples was 29.0 ± 19.1 ppm. Iodine content was below 30.0 ppm in 54.8% and below 15.0 ppm in 31.2% of salt samples. Mean per capita discretionary intake of household salt was 2.9 ± 1.8 g/day. Median UIC was 25.5 μg/L and Interquartile Range was 15.0 to 47.5 μg/L; 75.9% (221/291) of the children had UIC below 50.0 μg/L, indicating moderate status iodine nutrition. Median UIC was 34.3μg/L for children in households with salt, compared to 15.5 μg/L for children in households without salt, indicating severe iodine deficiency in the latter group. The three community groups had limited knowledge about importance of using iodised salt and consequences of iodine deficiency on health outcomes. This remote community has limited access to adequately iodised household salt due to high cost, inappropriate packaging, storage and food preparation, resulting in iodine deficiency. Strategies to increase iodine intake are needed.

## INTRODUCTION

Iodine is a trace element required for the biosynthesis of thyroid hormones, which are crucial to growth and development. Low bioavailability or deficiency of iodine may lead to a spectrum of disorders called iodine deficiency disorders (IDD) [1, 2]. This may include thyroid dysfunction with or without goitre, intellectual impairments, growth retardation, cretinism, increased pregnancy loss, and infant mortality. It is one of the most common causes of preventable impaired cognitive development [1, 2]. According to results of a meta-analysis [3] children living in areas with iodine deficiency had intelligence quotient (IQ) of 6.5–12.45 points lower than the IQ of those living in iodine-sufficient areas. Serious health consequences, such as cretinism and severe brain injury can be the manifestations of severe iodine deficiency [1-3]. Even mild iodine deficiency in pregnancy may lead to poorer cognitive outcomes in children, thus impairing their learning capacity and affecting the social and economic development of their countries [1, 4].These consequences of iodine deficiency are easily preventable by appropriate iodine intervention [1-3, 5]. The World Health Organisation (WHO), the United Nations Children’s Fund (UNICEF), and the International Council for the Control of Iodine Deficiency Disorders (ICCIDD) (now known as Iodine Global Network or IGN) have been assisting countries to implement appropriate strategies for the control and elimination of IDD worldwide [1, 2, 5]. Universal salt iodisation (USI), a policy of iodising all salt for human consumption, is the recommended strategy for the control and elimination of IDD in affected populations [1, 2, 5].

USI was implemented in Papua New Guinea (PNG) since June 1995 following promulgation of the PNG Salt Legislation, banning the importation and sale of non-iodised salt[6]. This was incorporated in the PNG Food Sanitation Regulation that was implemented in 2007 [7]. The effective implementation of USI requires systematic monitoring of the program [1].

The PNG National Nutrition Survey conducted in 2005 provided national data indicating that 92.5% of households with salt had adequately iodised salt and that the iodine status among non-pregnant women of child-bearing age was adequate – 170 μg/L[8]. The report further indicated that at the time of the survey 38% of households had no salt on the day of data collection and women in these households had lower iodine status than those in households with salt (114 μg/L and 203 μ respectively). In contrast to these national figures indicating adequate iodine status, a number of mini-surveys on iodine status among children and non-pregnant women carried out from 1998 to 2016 indicated prevalence of mild to moderate iodine deficiency in some districts in PNG, such as in school age children in Hella district, Southern Highland province; pregnant, lactating women and infants 0-4m in National Capital district; school age children in Aseki-Menyamya district, Morobe province and Lufa district, Eastern Highlands province [9-13]. Some of these areas are in mountainous regions, which may be more at risk of developing sub-optimal status of iodine nutrition [1-3, 5].

A recent study in the remote and mountainous area in Kotidanga LLG in Kerema district Gulf province in PNG reported moderate iodine status (median UIC 32.0 μg/L), stunting (57.6%), wasting (12.2%) and underweight (48.5%) among school-age children [14]. The authors indicated that more detailed study was urgently required to better understand the causes of the iodine status. The purpose of the current study was to therefore to assess in more detail the iodine status and the knowledge, attitudes and practice (KAP) relating to the use of iodised salt in this remote community. This was done by determination of the discretionary per capita intake of salt per day, the availability of adequately iodised salt in households, the iodine status of school children (age 6 –12 years) and the use of questionnaires to assess the KAP relating to the use of iodised salt.

## METHODS

### Study site and population

The study site was Kotidanga Rural LLG area, in Kerema district, Gulf province, PNG. Gulf province has rugged mountainous terrains, grassland flood plains and lowland river deltas. It shares land borders with six other provinces: Central, Western, Morobe, Simbu, Southern Highlands and Eastern Highlands[15]. There are two districts in Gulf province: Kerema and Kikori. The capital of Gulf province is Kerema, which is located in Kerema district. Kotidanga Rural LLG is also located in Kerema district. The local population is mainly the ‘Kamea’, which are members of the Angan-speaking tribal group [15]. The population of Kotidanga Rural LLG is 45,385 [16]. In Kotidanga Rural LLG, there is one hospital at Kanabea village, established by the Catholic mission in 1964; one Health Extension officer is servicing a population of around 20,000 Kamea people [16, 17]. Kotidanga Rural LLG is between 1200-1600 meters above sea level with temperatures between 12°C and 30°C and a yearly rainfall of 4000 – 7000 mm [18, 19]. There are no roads to the study site; the only way of getting there is by walking along mountain paths or by air transport. It usually takes about 3 days walking during daylight from the location of the hospital to the closest settlement in Kerema, Gulf province or Menyamya, Morobe province.

### Sample size

Calculation of sample size used a design effect of three, a relative precision of 10% and a confidence level (CL) of 95% [20]. As there was limited information on likely prevalence rates of iodine deficiency in the district, an assumed prevalence rate of 25% was used. With a predicted non-response rate of 10%, the sample size of 290 school-age children was obtained for the study area. This sample size was considered adequate for a mini-survey with limited resources and limited data across the study area.

### Study design and sampling

This was a prospective school and community based cross-sectional study carried out in Kotidanga Rural LLG in May-June 2017. There are 18 elementary, literacy and primary schools in Kotidanga Rural LLG. Simple random sampling was used to select nine literacy and primary schools in the study area. The total enrolments for each of the randomly selected schools including the ages of children in each of the grades were listed. Multistage cluster sampling was used to randomly select 291 school children in the age group 6 to 12 years.

### Collection of samples

The objectives of the study were explained to the head of each school and to the teachers, who were requested to communicate the information to the parents or caregivers. The data collectors visited the households of children. A teaspoon of salt was collected from households with salt at the time of the visit. Three brands of commercial salt were found to be available in the markets. A sample of each brand was purchased and analysed for iodine content.

### Discretionary intake of salt

To determine the discretionary intake of salt, sealed packets containing 250 g of iodised table salt were distributed to 30 randomly selected households of the 291 children. The number of individuals living in each household and eating food from the same cooking pot/hearth was counted and recorded. The head of the household was requested to use the salt as usual for cooking and eating. Each household was visited three days later to determine the amount of salt remaining in the packet. The number of individuals living in each household was again counted and recorded. The data obtained was used to estimate the average discretionary intake of salt per capita per day.

### Urinary iodine concentration (UIC)

For the determination of UIC, single urine samples were collected at the school from each of the 291 selected school children, after obtaining informed consent from their parents or caregivers. Each urine sample was kept in a properly labelled sterile plastic tube with a tight-fitting stopper that was further sealed with special plastic band.

Questionnaires knowledge, attitudes and practices (KAP) regarding use of iodised salt

The semi-structured FAO nutrition-related iodine deficiency questionnaire [21] was adapted for use in this study. It was pre-tested and then used to assess the KAP of randomly selected women visiting markets, market stallholders selling salt, and important stakeholders (community leaders, teachers, health staff) in Kotidanga Rural LLG.

The salt samples, urine samples and questionnaires were transported by airfreight to the Micronutrient Research Laboratory (MNRL) in the School of Medicine and Health Sciences (SMHS) University of Papua New Guinea (UPNG) for analysis.

### Exclusion criteria

Children below 6 years of age and above 12 years of age and those whose parents or caregivers did not give consent were excluded from the study.

### Analysis of salt and urine samples

The WYD Iodine Checker [22] was used for the quantitative assay of iodine content in salt collected from the households and purchased in the markets. Salt samples from the market were assayed 6 times and samples from households two to three times each, depending on the amount of salt collected. The Westgard Rules using Levy-Jennings Charts were used for internal bench quality control (QC) for daily routine monitoring of performance characteristics of the WYD Iodine Checker. The percent coefficient of variation (CV) ranges from 2.5% to 5.0% throughout the analysis.

The UIC was determined by Sandell-Kolthoff reaction after digesting the urine with ammonium persulfate in a water–bath at 100°C [1]. The Levy-Jennings Charts and the Westgard Rules were used for internal bench QC characterization of the assay method. The sensitivity (10.0 – 12.5 μg/L) and percentage recovery (95.0 ± 10.0%) of the urinary iodine (UI) assay were frequently used to assess the performance characteristics of the assay method. External QC monitoring of the assay procedure was by Ensuring the Quality of Urinary Iodine Procedures (EQUIP), which is the External Quality Assurance Program (QAP) of the Centers for Disease Control and Prevention (CDC), Atlanta, Georgia, USA.

### Data analysis and interpretation

The Statistical Package for Social Sciences (SPSS) software (version 17) and the Microsoft Excel Data Pack 2010 were used for statistical analyses of the data. Shapiro-Wilks test was used to assess normality of the data. Mann Whitney U and Wilcoxon W tests were used for differences between two groups; Kruskal-Wallis and Friedman were used for comparison of all groups. A p-value of< 0.05 was considered as statistically significant.

The criteria used for interpretation of the salt iodine data were based on the PNG salt legislation [6, 7]. According to the legislation all salt must be iodised with potassium iodate; the amount of iodine in table salt should be 40.0 to 70.0 ppm (mg/kg); the amount of iodine in other salt should be 30.0 to 50.0 ppm. These levels of iodine should be present at production or import level. WHO recommendations for iodine levels of food grade salt aim to provide 150μg iodine per day, assume 92% bioavailability, 30% losses from production to household level before consumption and variability of ±10% during iodisation procedures [5]. If 30% of iodine is lost from salt iodised per PNG food regulations, iodine content at household level should be at least 28 ppm (40 ppm minus 30%). This implies that in PNG the iodine content in salt in retail outlets or at the time of consumption should be at least 28.0 ppm [6, 7]. A cut-off of 30.0 ppm has been used in the analysis of this study by rounding up this figure. Global norms for iodine levels of salt at household level are 15 ppm based on the assumption that average salt consumption of 10g per day would provide the adult iodine requirement of 150 μg per day [1].

For the UIC data, the recommended WHO/UNICEF/ICCIDD [1] criteria were used to characterise the status of iodine nutrition among the school children. According to the criteria, a population of school-age children is considered iodine deficient if the median UIC is below 100.0 μg/L and more than 20% of the urine samples have UIC below 50.0 μg/L. The median UIC can also be used to indicate the severity of iodine deficiency; for example a population with median UIC <20.0 μg/L is considered severely deficient; moderately or mildly deficient if it is 20.0 to 49.0 μg/L or 50.0 to 99.0 μg/L respectively [1].

### Ethical approval

Ethical approval was obtained from the PNG National Department of Health Medical Research Advisory Committee (NDoH MRAC) and the Ethics and Research Grant committee in School of Medicine and Health Sciences (SMHS), University of Papua New Guinea (UPNG). Informed consent was also obtained from village authorities, and each adult participant and primary caretaker of the children.

## RESULTS

### Availability of salt in households

Of the 291 households selected a total of 289 households participated in this study, which gave a response rate of 99.3%. Salt was available in 186 out of 289 households (64.4%). All 186 salt samples were collected and analysed for their iodine content.

### Iodine content in salt from households and the markets

The mean (± STD) iodine content in salt from the households was 29.0 ± 19.1 ppm (mg/kg), the range was 1.3 to 58.2 ppm and the median was 30.5 ppm. The iodine content was below 30.0 ppm in 54.8% (102/186) of the salt samples, 31.2% (58/186) had iodine content below 15.0 ppm and 6.5% (12/186) had iodine content below 5.0 ppm.

Three brands of commercial salt were sold in the markets. One sample of each brand was purchased and coded as Brand A, Brand B and Brand C. The mean iodine content in brand A was 35.6 ± 6.0 ppm and median was 32.5 ppm. For brand B the mean was 40.0 ± 0.8 ppm and median was 36.5 ppm. Brand C, the mean was 46.9 ± 2.0 ppm and median was 43.6 ppm. Thus, the iodine content of samples of all three brands collected in the market was above 30.0 ppm as would be expected based on required iodine content at production/import level indicated in the PNG Food Sanitation Regulations for commercial salt [7].

### Discretionary per capita intake of salt and estimated per capita intake of iodine

The mean per capita discretionary intake of salt was 2.9 ± 1.8 g/day, with a range of 1.1 to 7.6 g/day and median of 2.3 g/day. The mean iodine content in the salt from the households was 29.0 ± 19.1 ppm. Thus, with a mean discretionary intake of 2.9 g of salt per capita per day, the calculated mean discretionary intake of iodine per capita per day was 84.1 ± 34.6 μg. Assuming that 20% of the iodine in salt is lost during storage and food preparation[1], the calculated per capita discretionary intake of iodine becomes 67.3 μg per day. This is below the 90.0 μg to 120.0 μg recommended daily requirement of iodine for school age children and 150 μg requirement for adults [1].

### Urinary Iodine Concentration (UIC)

Single urine sample was collected from each of the 291 school children; this gave a consent rate of 100%. The Shapiro-Wilks test for normality indicated that the frequency distribution curve of the UIC (μg/L) for all the children was not normally distributed (p = 0.001). This was confirmed by the Box-plot of the UIC data shown in Fig 1.

**Fig 1.**
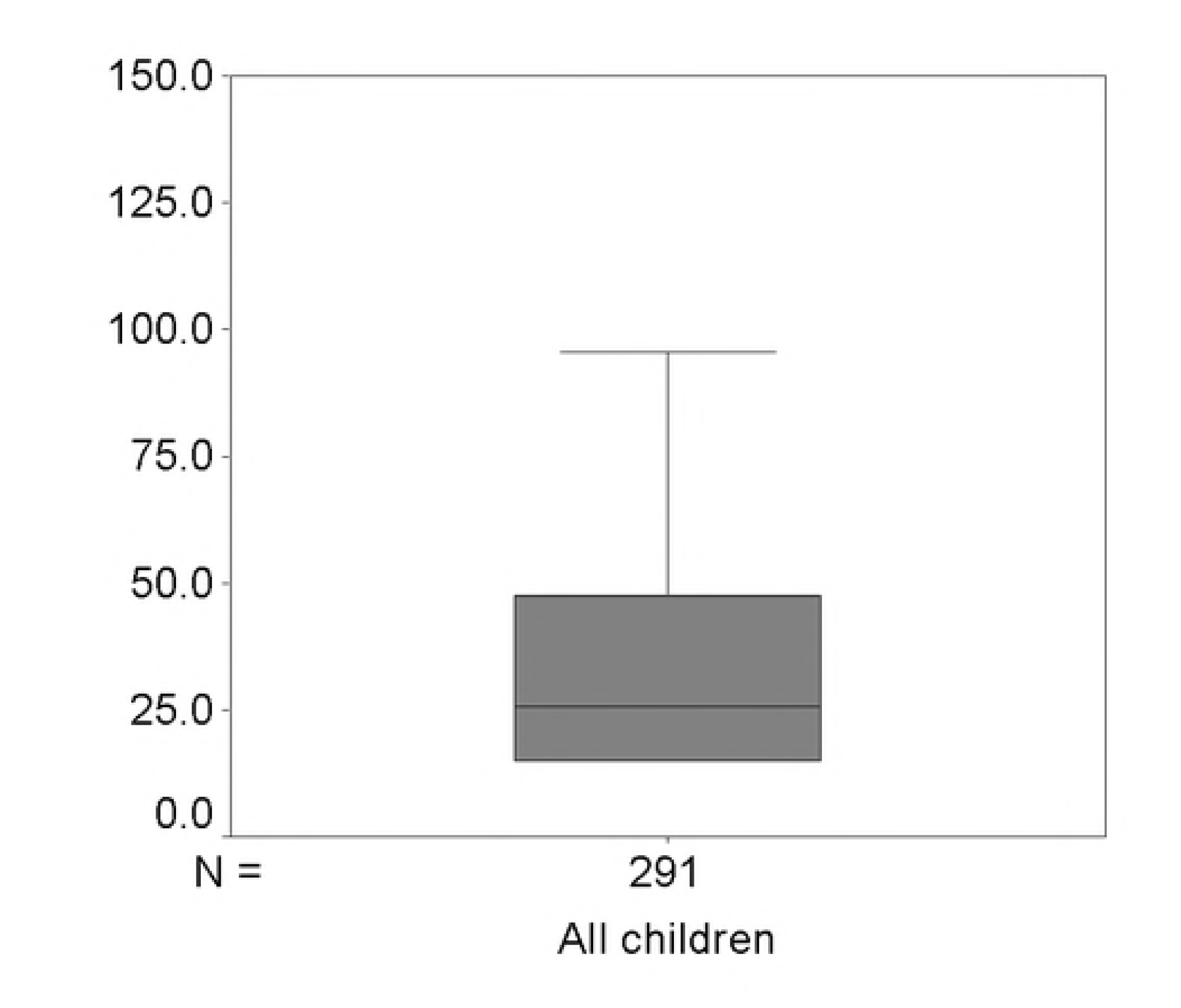
Box-plot of urinary iodine concentration (μg/L) for all the children.

The summary statistics of the UIC for the 291 children are presented in Table 1. The median UIC was 25.5μg/L and the Interquartile Range (IQR) was 15.0 to 47.5 μg/L. In addition, 75.9% (221/291) of the children had UIC below 50.0 μg/L. The median UIC of 25.5 μg/L indicates moderate iodine deficiency. It is notable that this is not far from <20.0 μg/L, which is the cut off for severely iodine deficient.

**Table 1:** Summary statistics of the urinary iodine concentration (μg/L) for all the children and for the male and female children.

| Parameters | All children | Male children | Female children |
| --- | --- | --- | --- |
| N | 291 | 175 | 114 |
| Median (µg/L) | 25.5 | 25.0 | 29.5 |
| Interquartile Range (IQR) (µg/L) | 15.0 – 47.5 | 15.0 – 51.0 | 15.0 – 46.5 |
| Percent (n) of children with UIC < 50.0 µg/L | 75.9% (221) | 80.0% (140) | 69.3% (79) |

The 291 UIC data for all the children were separated according to gender for further statistical analysis. The gender of 3 children was not indicated in the questionnaires. Those results were excluded from further analysis. Thus, of the 288 children, 175 (60.8%) were males and 113 (39.2%) were females. The Shapiro-Wilks test indicated that the UIC data for both male (p = 0.001) and female (p = 0.01) children were not normally distributed.

The median UIC for the male children was 25.0 μg/L and for the female children was 29.5 μg/L. Using the Mann-Whitney U and Wilcoxon W tests, no statistically significant difference (p = 0.264, 2-tailed) was indicated between the UIC of the male and female children. This was further confirmed by the Kruskal Wallis and Chi-Square tests (p = 0.254, 2-tailed). The UIC was below 50.0 μg/L in 80.0% (140/175) of male children and 69.3% (79/114) of the female children. The median UIC for both the male and female children indicated moderate iodine deficiency.

### Comparison of the UIC of children from households with and without salt

The 291 UIC results were separated based on the availability of salt in the households of the children on the day of the visit. Of the 291 households selected a total of 289 households participated in this study, thus 289 UIC results were used for the statistical analysis. Of the 289 households salt was collected in 186 (64.4%), but not available in 103 (35.6%) of the households.

The Box-plots of the UIC data for the two groups are presented in Fig 2; they indicate that the UIC data were not normally distributed, which was supported by the Shapiro-Wilks test for normality of distribution (p = 0.01 and P = 0.001). The summary statistics of the UIC data for both cases are presented in Table 2. The median UIC was 34.3 μg/L for children in households with salt, indicating moderate iodine deficiency; compared to 15.5 μg/L for children in households without salt, indicating severe iodine deficiency. Statistically significant difference (p = 0.02; 2-tailed) was indicated between the UIC of children in households with salt compared to those in households without salt. The UIC was below 50.0 μg/L in 72.0% (134/186) of children in households with salt compared to 82.5% (85/103) of children in households without salt.

**Fig 2.**
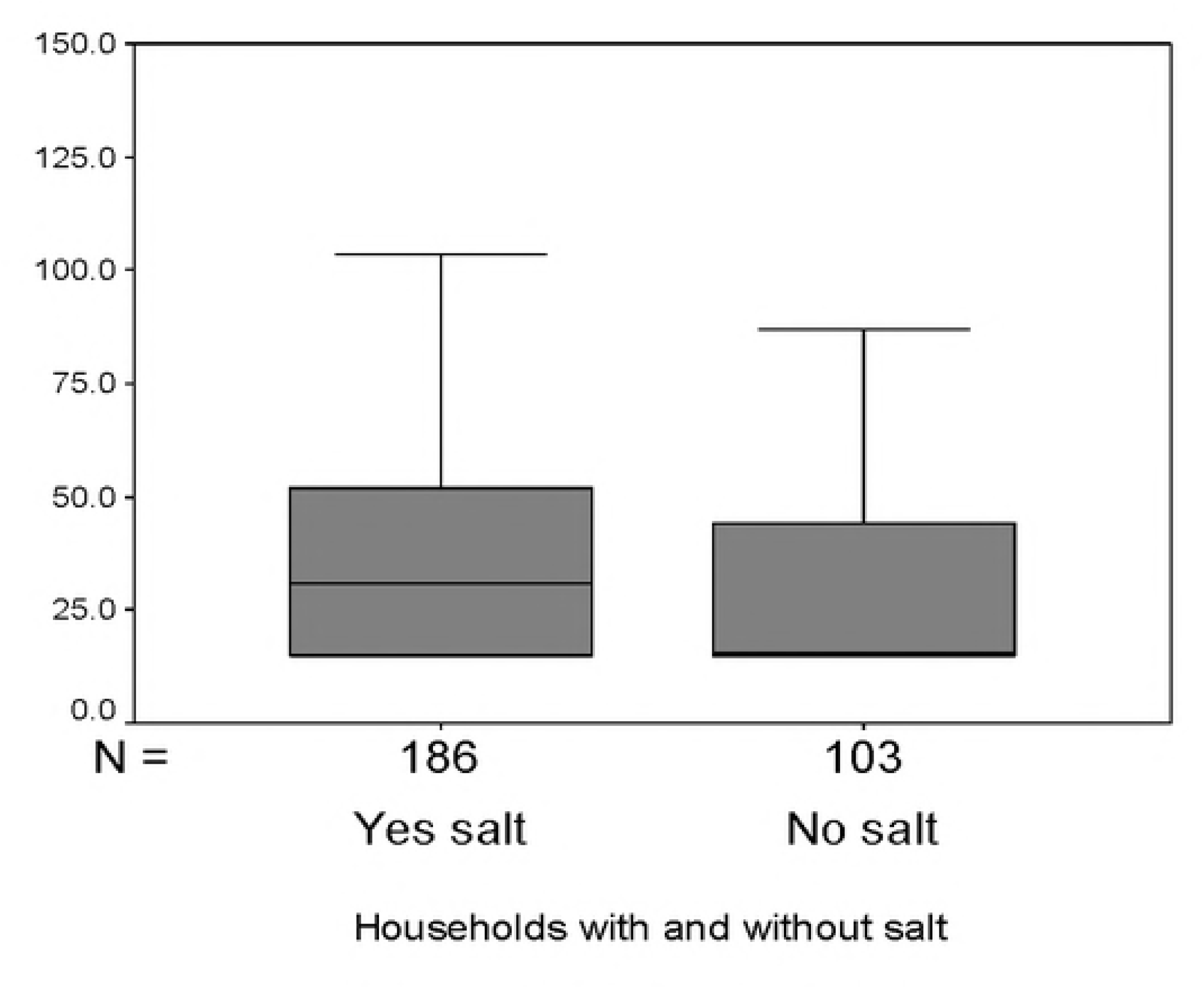
Box-plots of urinary iodine concentrations (μg/L) for children in the households with and without salt.

**Table 2.** Summary statistics of urinary iodine concentration of children in households with salt and without salt.

|  | Availability of iodised salt in households |  |
| --- | --- | --- |
|  | YES | NO |
| N | 186 (64.4%) | 103 (35.6%) |
| Median UIC (µg/L) | 34.3 | 15.5 |
| IQR (µg/L) | 15.0 – 52.0 | 15.0 – 44.3 |
| Percent (n) of children with UIC< 50 µg/L | 72.0% (134) | 82.5% (85) |

### Assessment of knowledge, attitudes and practices (KAP)

All five markets across the study area were visited. Women visiting the markets to purchase items and stallholders selling salt in the markets were randomly selected to participate in this section of the study. In addition, important community stakeholders in their place of work (community leaders, teachers, and health staff) were also selected. The variation in the number of participants in the three groups was due to logistical reasons.

### Women visiting markets

A total of 153 women completed the questionnaires. The mean age of the women was 32.3 ± 8.2 years, the range was 18.0 to 50.0 years and the median age was 32.0 years.

The questionnaire results are presented in Table 3. The results show that 91% of the women cannot read, 87% did not listen to radio and 97% indicated that they do not work for money. In response to questions on the use of salt, 47% reported to always use salt at home, 49% reported they do not always use salt at home and only buy salt when they have money and 4% reported to use only traditional salt. Of the 47% that always use salt at home, 57% reported they use salt for cooking only and 43% use it for cooking and also adding to food before eating. When asked about iodised salt, 15.5% indicated to have some knowledge and 84.5% did not have any knowledge about iodised salt. Furthermore, 65% of the women reported they store salt in bamboo stem covered with a leaf and 35% store salt in the original plastic bag. A total of 86% said it is good to prepare food with salt, 14% were unsure about salt in food; 88% indicated that it is difficult to buy salt because they usually do not have money and 5% indicated salt is not available.

**Table 3.**
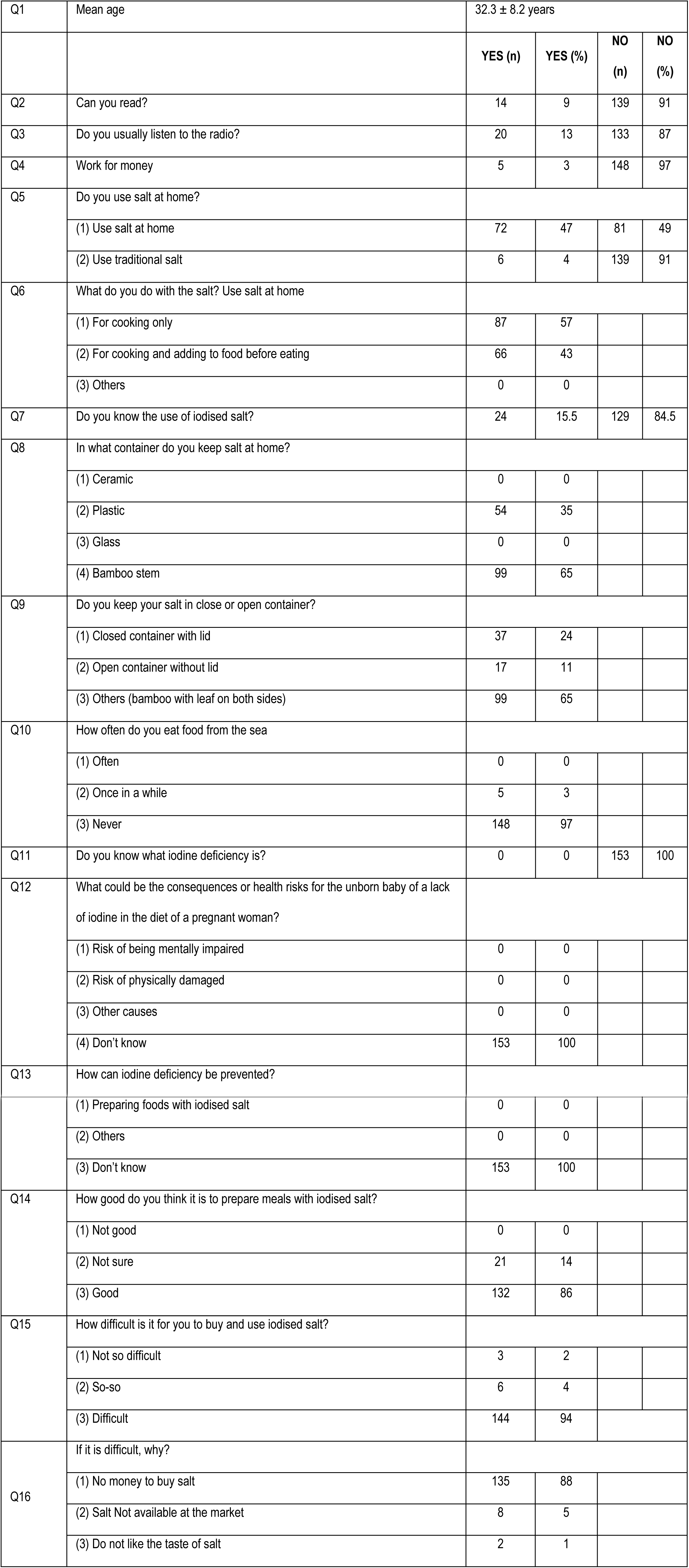
Knowledge, attitude, practice women visiting markets (n=153).

|  |  |  |  |  |  |
| --- | --- | --- | --- | --- | --- |
| Q1 | Mean age | 32.3 ± 8.2 years |  |  |  |
|  |  | YES (n) | YES (%) | NO (n) | NO (%) |
| Q2 | Can you read? | 14 | 9 | 139 | 91 |
| Q3 | Do you usually listen to the radio? | 20 | 13 | 133 | 87 |
| Q4 | Work for money | 5 | 3 | 148 | 97 |
| Q5 | Do you use salt at home? |  |  |  |  |
|  | (1) Use salt at home | 72 | 47 | 81 | 49 |
|  | (2) Use traditional salt | 6 | 4 | 139 | 91 |
| Q6 | What do you do with the salt? Use salt at home |  |  |  |  |
|  | (1) For cooking only | 87 | 57 |  |  |
|  | (2) For cooking and adding to food before eating | 66 | 43 |  |  |
|  | (3) Others | 0 | 0 |  |  |
| Q7 | Do you know the use of iodised salt? | 24 | 15.5 | 129 | 84.5 |
| Q8 | In what container do you keep salt at home? |  |  |  |  |
|  | (1) Ceramic | 0 | 0 |  |  |
|  | (2) Plastic | 54 | 35 |  |  |
|  | (3) Glass | 0 | 0 |  |  |
|  | (4) Bamboo stem | 99 | 65 |  |  |
| Q9 | Do you keep your salt in close or open container? |  |  |  |  |
|  | (1) Closed container with lid | 37 | 24 |  |  |
|  | (2) Open container without lid | 17 | 11 |  |  |
|  | (3) Others (bamboo with leaf on both sides) | 99 | 65 |  |  |
| Q10 | How often do you eat food from the sea |  |  |  |  |
|  | (1) Often | 0 | 0 |  |  |
|  | (2) Once in a while | 5 | 3 |  |  |
|  | (3) Never | 148 | 97 |  |  |
| Q11 | Do you know what iodine deficiency is? | 0 | 0 | 153 | 100 |
| Q12 | What could be the consequences or health risks for the unborn baby of a lack of iodine in the diet of a pregnant woman? |  |  |  |  |
|  | (1) Risk of being mentally impaired | 0 | 0 |  |  |
|  | (2) Risk of physically damaged | 0 | 0 |  |  |
|  | (3) Other causes | 0 | 0 |  |  |
|  | (4) Don't know | 153 | 100 |  |  |
| Q13 | How can iodine deficiency be prevented? |  |  |  |  |

|  |  |  |  |
| --- | --- | --- | --- |
|  | (1) Preparing foods with iodised salt | 0 | 0 |
|  | (2) Others | 0 | 0 |
|  | (3) Don't know | 153 | 100 |
| Q14 | How good do you think it is to prepare meals with iodised salt? |  |  |
|  | (1) Not good | 0 | 0 |
|  | (2) Not sure | 21 | 14 |
|  | (3) Good | 132 | 86 |
| Q15 | How difficult is it for you to buy and use iodised salt? |  |  |
|  | (1) Not so difficult | 3 | 2 |
|  | (2) So-so | 6 | 4 |
|  | (3) Difficult | 144 | 94 |
| Q16 | If it is difficult, why? |  |  |
|  | (1) No money to buy salt | 135 | 88 |
|  | (2) Salt Not available at the market | 8 | 5 |
|  | (3) Do not like the taste of salt | 2 | 1 |

### Stallholders selling salt

Gender distribution of the 36 stallholders selling salt that participated showed 23 (64%) were male and 13 (36%) were female. The combined mean age was 30.4 ± 5.6 years, the range was 22 to 45 years and the median age was 30.0 years.

The questionnaire results are presented in Table 4. Among the stallholders 67% reported that cannot read and 67% do not listen to the radio. All stallholders (100%) reported to only sell iodised salt. All salt brands sold at markets indicated “iodised salt”, however some salt had been repackaged into smaller quantities by the stallholder and it was unclear what brand of salt this was. The mean monthly sale of salt (250g packets) across the five markets was 22.5 packets. Brand A salt was sold by 92% stallholders, 6% sold brand B and only 2% sold brand C. Each stallholder sold only one brand of salt. Of the 36 stallholders 44% had some knowledge about iodised salt, although all of them (100%) had no knowledge about iodine deficiency and its potential consequences or health risks for the population. However, 88% indicated that it is good to prepare food with salt. In addition, 96% of the stallholders indicated that it is difficult to bring salt to Kotidanga LLG to sell because it takes about a minimum of 3 days to walk (one-way) to major settlements including Lae, Menyamya, Kerema or Port Moresby to purchase the salt. Furthermore, 89% of stallholders indicated that people do not buy salt because they do not have money to buy it and 11% indicated that people do not like the taste of salt.

**Table 4.** Knowledge, attitude and practice of stallholders selling salt (n = 36).

|  |  |  |  |  |  |
| --- | --- | --- | --- | --- | --- |
| Q1 | Gender | Males 64%; Females 36% |  |  |  |
| Q2 | Combined Mean age | 30.4 ± 5.6 years |  |  |  |
|  |  | YES (n) | YES (%) | NO (n) | NO (%) |
| Q3 | Can you read? | 12 | 33 | 24 | 67 |
| Q4 | Do you usually listen to the radio? | 12 | 33 | 24 | 67 |
| Q5 | What salt do you sell at the stall? |  |  |  |  |
|  | (1) Iodised salt | 36 | 100 | 0 | 0 |
|  | (2) Not iodised salt | 0 | 0 | 36 | 100 |
| Q6 | How many packets of salt do you sell each month?<br>(Mean monthly sale of packets (250g) of salt): | 22.5 packets per month |  |  |  |
| Q7 | What brands of salt do you sell? |  |  |  |  |
|  | Brand A: | 33 | 92 |  |  |
|  | Brand B: | 2 | 6 |  |  |
|  | Brand C: | 1 | 2 |  |  |
|  | Other brands | 0 | 0 |  |  |
| Q8 | Knowledge of iodised salt | 16 | 44 | 20 | 56 |
| Q9 | What are reasons why people are not buying iodised salt? |  |  |  |  |
|  | (1) Lack of money | 32 | 89 |  |  |
|  | (2) They don't like the taste | 4 | 11 |  |  |
|  | (3) Other reasons | 0 | 0 |  |  |
| Q10 | Do you know what iodine deficiency is? | 0 | 0 | 36 | 100 |
| Q11 | What could be the consequences or health risks for the unborn baby of a lack of iodine in the diet of a pregnant woman? |  |  |  |  |
|  | (1) Risk of being mentally impaired | 0 | 0 |  |  |
|  | (2) Risk of physically damaged | 0 | 0 |  |  |
|  | (3) Other reasons | 0 | 0 |  |  |
|  | (4) Don't know | 36 | 100 |  |  |
| Q12 | How can iodine deficiency be prevented? |  |  |  |  |
|  | (1) Eat / prepare foods with iodised salt | 0 | 0 |  |  |
|  | (2) Other reasons | 0 | 0 |  |  |
|  | (3) Don't know | 36 | 100 |  |  |
| Q13 | How good do you think it is for people to prepare meals with iodised salt? |  |  |  |  |
|  | (1) Not good | 0 | 0 |  |  |
|  | (2) Not sure | 4 | 12 |  |  |
|  | (3) Good | 32 | 88 |  |  |
| Q14 | How difficult is it for you to buy iodised salt in Port Moresby, Kerema, Lae and Menyamya? |  |  |  |  |
|  | (1) Not so difficult | 0 | 0 |  |  |
|  | (2) So-so | 1 | 4 |  |  |
|  | (3) Difficult | 35 | 96 |  |  |
| Q15 | If it is difficult, why? |  |  |  |  |
|  | Major settlement to purchase salt is at minimum 3 days walking distance | 35 | 96 |  |  |

### Stakeholders in the community

A total of 43 important stakeholders (community leaders, teachers, and health staff) from 9 villages across Kotidanga Rural LLG participated in this section. When separated according to gender, 33 (77%) were males and 10 (23%) were females. Their combined mean age was 35.8 ± 6.4 years, the range was 26 to 57 years and the median age was 36.0 years.

Table 5 shows the results from the questionnaires. All the stakeholders (100%) reported that they can read. Majority (74%) reported that do not listen to the radio. Only 14% had knowledge about the use of iodised salt; 5% indicated that it is best to store iodised salt in a closed container at home; all of them (100%) indicated that people do not buy salt because of lack of money to purchase it. Furthermore, 92% responded that they do not have any knowledge about iodine deficiency; 89% do not know how iodine deficiency could be prevented, but 11% stated that they were using iodised salt and 72% indicated that it is good to prepare food with iodised salt. All (100%) indicated that it is difficult for community members to purchase salt mainly because they do not have money.

**Table 5.** Knowledge, attitude and practice of stallholders (health, education staff, community leaders) (n = 43).

|  |  |  |  |  |  |
| --- | --- | --- | --- | --- | --- |
| Q1 | Gender | Males 77%; Females 23% |  |  |  |
| Q2 | Combined mean age | 35.8 years |  |  |  |
|  |  | **YES**<br>**(n)** | **YES**<br>**(%)** | **NO (n)** | **NO (%)** |
| Q3 | Can read | 43 | 100 |  |  |
| Q4 | Listen to radio | 11 | 26 | 32 | 74 |
| Q5 | Do you know the use of iodised salt? | 6 | 14 | 37 | 86 |
| Q6 | Are you aware how iodised salt needs to be stored at home? | 2 | 5 | 41 | 95 |
| Q7 | Why are people not buying iodised salt? |  |  |  |  |
|  | (1) Lack of money | 43 | 100 |  |  |
|  | (2) They don't like the taste | 0 | 0 |  |  |
|  | (3) Other reasons | 0 | 0 |  |  |
| Q8 | Do you know what iodine deficiency is? | 3 | 8 | 40 | 92 |
| Q9 | What could be the consequences or health risks for the unborn baby of a lack of iodine in the diet of a pregnant woman? |  |  |  |  |
|  | (1) Risk of being mentally impaired | 3 | 6 |  |  |
|  | (2) Risk of being physically damaged | 1 | 2 |  |  |
|  | (3) Other reasons | 0 | 0 |  |  |
|  | (4) Don't know | 39 | 92 |  |  |
| Q10 | How can iodine deficiency be prevented? |  |  |  |  |
|  | (1) Eat / prepare foods with iodised salt | 5 | 11 |  |  |
|  | (2) Other reasons | 0 | 0 |  |  |
|  | (3) Don't know | 38 | 89 |  |  |
| Q11 | Are you aware of foods rich in iodine? | 3 | 6 | 40 | 94 |
| Q12 | How likely is it, do you think that children in this community lack iodine? |  |  |  |  |
|  | (1) Not likely | 0 | 0 |  |  |
|  | (2) Not sure | 43 | 100 |  |  |
|  | (3) Not likely | 0 | 0 |  |  |
| Q13 | How serious do you think is a lack of iodine in the body? |  |  |  |  |
|  | (1) Not serious | 0 | 0 |  |  |
|  | (2) Not sure | 43 | 100 |  |  |
|  | (3) Serious | 0 | 0 |  |  |
| Q14 | How do you think it is for people to prepare meals with iodised salt? |  |  |  |  |
|  | (1) Not good | 0 | 0 |  |  |
|  | (2) Not sure | 12 | 28 |  |  |
|  | (3) Good | 31 | 72 |  |  |
| Q15 | How difficult is it for community members to buy iodised salt? |  |  |  |  |
|  | (1) Not so difficult | 0 | 0 |  |  |
|  | (2) So-so | 0 | 0 |  |  |
|  | (3) Difficult | 43 | 100 |  |  |
| Q16 | If difficult, why? |  |  |  |  |
|  | (1) Market does not sell salt | 0 | 0 |  |  |
|  | (2) People do not like the taste | 0 | 0 |  |  |
|  | (3) People have no money to buy salt | 43 | 100 |  |  |
|  | (4) Other reasons | 0 | 0 |  |  |

## DISCUSSION

According to recent WHO guidelines, all food-grade salt, used in household and food processing, should be fortified with iodine as a safe and effective strategy for the prevention and control of IDD in populations living in stable and emergency settings [5]. Salt is considered an appropriate vehicle for fortification with iodine because it is widely consumed by virtually all population groups in all countries with little seasonal variation in consumption patterns [5]. Thus the percentage of household salt with iodine content between 15.0 and 40.0 ppm in a representative sample of households must be equal to or greater than 90% [1].

A basic premise of the WHO global strategy is that salt is consumed by all population groups. However, on the day of the visit, salt was available for collection in only 64.4% (186/289) of the households in the study area. One of the major reasons why salt was not available in 35.6% (103/289) of the households appears to be because it is considered not affordable and only limited amounts are occasionally purchased.

Of the 64.4% of households which did have salt on the day of the survey, only 45.2% (84/186) had salt with iodine content ≥ 30.0 ppm cut-off point as expected according to the PNG Salt Legislation for adequately iodised salt. However, 68.8% (128/186) of households had salt with iodine content ≥ 15.0 ppm, the WHO/IGN/UNICEF global cut-off for adequately iodised salt at household level [1]. Thus, in the present study the percentage of total households that had adequately iodised salt with iodine content ≥ 30 ppm was only 44.3% (128/289). This is significantly lower than the WHO/IGN/UNICEF recommended 90% coverage that should indicate effective implementation for iodised salt in the household [1].

The 44.3% of households obtained in this study was lower than those in other studies in various districts in PNG that reported higher percentage of households with adequately iodised salt; 95.0% in Hella District in 2004 [9], 94.5% in National Capital district in 2006 [23], 95.0% in National Capital district in 2009 [11], 78.0% in Morobe and Eastern Highlands Provinces in 2013 [12]; and 66% in Karimui-Nomane district. The results were higher compared to 28% in Sina Sina Yonggomugl district, Simbu Province in 2017 [24].

A further limitation regarding the impact of salt iodisation in PNG is the apparent low consumption of discretionary salt. In 2010, a systematic analysis of 24-hour urine sodium excretion and dietary surveys worldwide estimated the total salt consumption, including non-discretionary sources of salt. The study found salt intake to vary widely between countries – from 3.76 g/day in Kenya to 14.3 g/day in Uzbekistan with a global average of 10.6 g/day [25]. The 2.9 ± 1.8 g mean daily discretionary intake of salt per capita obtained in this study was well below the lowest intake indicated in the study [25] and the 6.23 g/day estimated for PNG [25]. The present study assessed only the discretionary salt intake, which did not include, for example salt contained in processed foods such as bouillon cubes or instant noodles that are not commonly used for food preparation in this community. The 2.9 ± 1.8 g is also lower than the values obtained in other studies in PNG that also assessed only discretionary salt intake by the same procedure; 6.6 g in Lae city [13], 4.7 g in Morobe [12] and 5.0 g in Simbu province [24]. It was also below the 10.0 g per capita per day salt intake used in formulating the PNG standards for iodine content in salt indicated in the PNG Salt Legislation [6, 7].

The low per capita discretionary intake of salt and the low iodine content in the salt from a proportion of the households resulted in the calculated low iodine (67.28 μg per day) per capita intake of iodine obtained in the present study in the households with salt. This can lead to iodine deficiency among the vulnerable groups in the community and is further exacerbated by the low access to commercial salt recorded in this study.

This study found that the mean and median iodine content of salt in the households were lower than the salt samples collected in the market. Recognising that 92% of stallholders sold Brand A (mean iodine content = 35.6 ppm), 6% sold Brand B (mean = 40.8 ppm) and 2 % sold Brand C (mean = 46.9 ppm) a weighted average content of all salt at market level is 36.1 ppm compared to that at household level of 29.0 ppm. This suggests some losses of iodine between the market and households. Losses of iodine may occur because some stallholders repacked salt from the 250 g original packages into smaller amounts, in plastic bags or wrapped in paper or leaves and also because most households (65%) reported storing salt uncovered in the home, close to the fire. These practices may result in significant loss of iodine as reported by several authors [26-31]. According to these authors, the loss of iodine in salt can occur under various conditions including the type of retail packaging using unconventional packets, storage and transport at temperatures above 30°C and the high relative humidity. Furthermore, some iodine may be lost during preparation and cooking of meals; more than half of the women (57%) reported adding salt to water for cooking only to then discard the liquid bearing much of the salt.

A previous survey in 2015 in the same area, recorded a median UIC of 32 μg/L in 151 school age children [14]. In this survey with data collected during May-June 2015, the median UIC of 25.5 μg/L of 291 school age children suggest a further deterioration in iodine status in this vulnerable population group. No data on access to adequately iodised salt was collected in the last survey; only data on use of commercial salt was collected. The statistically significant difference in the median UIC among children in households with salt (34.3 μg/L) and those in households without salt (15.5 μg/L), the estimate of discretionary salt intake and iodine levels of salt in households and markets in the present study confirm that the poor iodine status among the children is likely related to low access to commercial salt, low discretionary salt intake and inadequate iodine levels in a proportion of the limited amount of salt being consumed.

Similar situations have been recorded in some rural and remote communities in PNG, in Hella district, Southern Highland province; in Aseki-Menyamya district, Morobe province and Lufa district, Eastern Highlands province, Karimui-Nomane and Sina-Sina Yonggomugl districts in Simbu province [9, 11, 24]. While nationally iodine status may be adequate due to access to adequately iodised salt [8], there appear to be pockets of remote communities that have low access to commercial salt and are at risk of iodine deficiency.

The results of the KAP indicated that most of the respondents in the three groups had very limited knowledge about the importance and benefits of using iodised salt and also the consequences of iodine deficiency on the health of the population. The regular diet in this remote community living at high-altitude and with high precipitation is low in natural sources of iodine; they rarely consumed fish or other sea-foods, thus risk of iodine deficiency is high without interventions to increase iodine intake [1]. Access to commercial salt was found to be very low in this community due to insufficient disposable income and remoteness, which drives up the cost of external commodities, such as salt. Poor repacking and household storage and food preparation practices may also have caused losses of iodine from the small amount of salt consumed.

It is important to acknowledge the questionnaire used may have limitations. This questionnaire was based on a FAO questionnaire validated for use in developing countries [21]. However, responses on the questions “do you use salt at home” and “do you use traditional salt” and “reasons why people are not buying iodised salt” may not reflect all the response options or are mutually exclusive.

While lack of finance is one of the major barriers for using iodised salt regularly in the households, inappropriate transport, storage and usage practices, both by stallholders and in the households are major contributing factors that may reduce the iodine content in the salt. Important strategies, such as, increasing advocacy and improving “health literacy” on the importance of consumption of iodised salt should be carried out among key target groups in the community [32].

It is important to carry out intensive nutrition education and information provision, together with an awareness raising campaign, to advocate for changes in the practices among stallholders and in the households that can affect the iodine content in salt and also to improve regular intake of iodised salt. Any education, promotion and communication of the importance of improving iodine nutrition should be tailored to the low literacy levels of this community. Social mobilisers should carryout regular visits to the households for face-face discussion on the use of iodised salt, using appropriate visual resources and tools. The importance of iodised salt should be included in health education of children in schools, and in discussions in churches and markets. Similar approaches have been carried out in remote communities in other countries with reasonable success [29, 33-36].

A potential strategy to increase access to commercial (iodised) salt in this and other similar communities is to seek opportunities to increase its availability, such as by subsidising transport costs or improving distribution networks. Alternative or complementary strategies to increase iodine intake should also be explored such as the fortification of alternative food vehicles that are more readily available, such as cereal grains including rice or wheat flour, edible vegetable oil, or condiments and seasonings [37], or targeted distribution of iodine or combined micro-nutrient supplements to vulnerable groups in the community, in particular reproductive age women [38].

## CONCLUSIONS

The percentage of households with adequately iodised salt was significantly lower than the recommended coverage that should indicate effective implementation of the USI strategy at the household level. The low median UIC indicates moderate status of iodine deficiency and insufficient intake of iodine among the school age children. Most of the respondents in the three groups had very limited knowledge about the importance and benefits of using iodised salt and also the consequences of iodine deficiency on the health of the population. Lack of salt iodisation is not a major risk factor for the development of iodine deficiency among the population in the Kotidanga Rural LLG, because the commercial salt sold by the stallholders in the markets was adequately iodised. Issues in accessing remote markets prompt prohibitive increases in retail salt prices and inappropriate storage and packaging practices by local traders; improper storage and food preparation practices in households may have caused loss of iodine in the salt. The combination of these factors ultimately resulted in the low availability of adequately iodised salt in the households.

A specially designed intervention strategy is recommended to address the fundamental problems relating to iodine deficiency and insufficient iodine intake in the study area. This strategy would include ensuring effective low-cost transportation of iodised salt to the remote communities; reducing the cost of salt by producing smaller (25 to 50 g) quantities of commercial iodised salt in appropriate packaging material; appropriate storage methods in households; facilitation of training for community champions on the importance of iodised salt and the subsequent implementation of iodine nutrition education and promotion in the community and school settings. Essential for successful iodine nutrition intervention is government commitment, clear policy and program direction and ongoing monitoring to ensure full compliance and sustained impact.

## ACKNOWLEDGEMENTS

The authors thank the Kotidanga LLG community, schools, students and health staff for their participation in this study. We acknowledge the support of the Chief Technical Officer and other technical staff members in the Division of Basic Medical Sciences, School of Medicine and Health Sciences, University of PNG.

The findings and conclusions in this manuscript are those of the authors; they do not represent the official position of the institutions and authors’ organisations. The authors declare no conflicts of interest and they did not receive any research funding for this project. The analyses of iodine in salt and urine were carried out in the Micronutrient Research Laboratory, School of Medicine and Health Sciences, University of Papua New Guinea.

## Author contributions

Conceptualisation: Janny M Goris, Victor J Temple

Formal analysis: Victor J Temple, Janny M Goris, Nienke Zomerdijk Investigation/field work: Janny M Goris

Methodology: Victor J Temple, Janny M Goris

Project administration: Janny M Goris, Victor J Temple

Resources: Victor J Temple, Janny M Goris

Validation: Victor J Temple

Visualisation: Victor J Temple

Writing: Janny M Goris, Victor J Temple, Nienke Zomerdijk

Writing-review and editing: Janny M Goris, Victor J Temple, Karen Codling, Nienke Zomerdijk

